# Ventral-to-dorsal electrocyte development in electric organs of electric eel (*Electrophorus*)

**DOI:** 10.1101/2024.08.21.606117

**Authors:** Sinlapachai Senarat, Ayako Matsumoto, Tatsuki Nagasawa, Shintaro Sakaki, Daichi Tsuzuki, Kazuko Uchida, Makoto Kuwahara, Masato Nikaido, Eiichi Hondo, Atsuo Iida

## Abstract

Electric eels (*Electrophorus*) are renowned for their ability to generate electric discharge, which is used for prey capture and defense. Their electric organs (EOs) are located along the lateral–ventral region of the tail and contain electrocytes, which are multinucleated syncytium cells. The developmental origin of the electrocyte is mesodermal lineage cells observed in the ventral part of the myotome. However, it is unclear whether these precursor cells are also maintained in later stages and contribute to electric organ growth in adulthood. In this study, we report regional differences in cell morphology within the main EO (mEO) and identify a candidate cell population for electrocyte progenitors at the juvenile stages of Electrophorus. The cell morphology and distribution from the ventral terminal to the dorsal region of the mEO suggest the segregation of progenitors from the ventral cluster and their gradual transformation into mature multinucleated electrocytes via cell fusion and layering along the dorsal axis. Immunohistochemistry revealed strong expression of sodium-potassium adenosine triphosphatase (Na^+^/K^+^-ATPase), a key component in generating electric discharge in the mEO, across most mEO regions, except in the ventral cluster cells. Based on these observations, we propose that electrocyte progenitors develop from ventral cluster cells in the mEO and differentiate into mature multinucleated cells as they migrate dorsally. This is the first report to approach the developmental process of *Electrophorus* electrocytes from cell morphology and genetic profiles. Our findings represent a breakthrough in understanding the differentiation of electrocytes during the growth stages of *Electrophorus*.

## Introduction

Electric fish include multiple classified species, such as electric rays, catfish, and eels. As Charles Darwin described in his work *On the Origin of Species* (Chapter 6: Difficulties on Theory), “The electric organs of fishes offer another case of special difficulty; for it is impossible to conceive by what steps these wondrous organs have been produced” (Darwin, 1859). Darwin’s question of how these electric species appeared independently in each taxon is difficult and remains elusive. Here, we address this question from the perspective of cell and molecular biology using electric eels (*Electrophorus*).

The electric eel, an electric fish, was originally described by Carl Linnaeus in 1766 as *Gymnotus electricus* and was later revised as *Electrophorus electricus* by Theodore Gill in 1864 (Jordan, 1963). Recently, they have been reclassified into three distinct species, *E. electricus*, *Electrophorus varii*, and *Electrophorus voltai*, based on both morphological and genetic sequence data (de Santana et al., 2019). Electric eels are known for their ability to generate electric discharges, with a maximum capacity of 860 V for *E. voltai* (de Santana et al., 2019). They possess three types of electric organs (EOs): the main electric organ (mEO), Hunter’s organ, and Sachs’ organ, which are located in the lateral-ventral region of the tail (Williamson, 1775; Hunter, 1775; Sachs, 1877). The first electric organ, called the larval organ, appears in the ventral trunk region as early as the juvenile stage (total length [TL] 15 mm) (Schwassmann et al., 2014). Most larval organs grow into the mEO, but the caudal region is transformed into the Sachs’ organ. The Hunter’s organ has a different origin, the anal fin organ. The differentiation of the three organs is completed at approximately a TL of 23 cm (Moller, 1995). The mEO is predicted to play a role in high-voltage discharge for prey capture or defense, whereas the other organs are considered to play a role in low-voltage discharge for sensing and communication (Xu et al., 2021). The cells responsible for electric discharge in an EO are multinucleated syncytia called electrocytes (Luft, 1957). Early studies suggested that electrocytes are derived from muscle lineage precursors (Fritsch, 1883). Recent histological observations in juvenile *Electrophorus* suggest that the larval organ consists of electrocytes budded from a specified cell cluster named the electromatrix, located in the ventral region of the myotome (Schwassmann et al., 2014). More detailed observations and multifaceted analyses of *Electrophorus* embryos are required to determine the distinct origin and developmental mechanisms of the first electrocytes. However, artificial breeding methods have not yet been established; therefore, obtaining embryos in the laboratory is challenging. As an alternative model, the weakly electric fish, *Brachyhypopomus gauderio*, a species related to *Electrophorus* belonging to the same order (Gymnotiformes), was used to observe developmental stages, including EOs, under laboratory conditions (Alshami et al., 2020). In this species, cell clusters consisting of Pax7^+^/Myosin^-^ cells in the ventral region of the myotome were reported as the EO primordium. Based on these studies, the developmental origin of electrocytes in electric species of Gymnotiformes is considered to be the ventral cell cluster derived from somite lineage cells rather than from differentiated myocytes. In these studies, electrocyte lineage cells were classified into two populations: mature electrocytes and immature cells located in the ventral cluster. However, either or both populations may be classified into further substages. In this case, their clarification is expected to lead to a more detailed description of the differentiation process of electrocyte lineage cells. Moreover, researchers have focused on the developmental origin of electrocytes; however, the source of these cells during growth remains elusive.

From another perspective, advances in molecular biology methods have enabled the comparison of gene expression between EOs and other tissues, including skeletal muscle. In EOs, some muscle characteristic genes were downregulated, whereas other genes not expressed in the muscle were upregulated (Gallant et al., 2014; Gallant, 2019). Specifically, genes for ion transporters or channels, such as *atp1a*, *chrna*, *scn4aa*, and *kcnj12*, are upregulated not only in electric eels, but also in other electric fish species (Thompson et al., 2018; LaPotin et al., 2022). Some functional mutations are found in the genes encoding the ion transporters of *Electrophorus* compared to non-electric species. Substitutions in the amino acid sequences of these proteins have been predicted to contribute to high-voltage discharge in electrocytes (Traeger et al., 2017), and this modification is considered crucial for high-voltage electric discharge in *Electrophorus* spp. The molecular genetic characteristics of mature electrocytes are well known; however, how these expressed genes affect the differentiation of electrocytes remains unknown. Here, we report the progenitors of electrocytes based on histological observations and immunohistochemistry. Understanding the spatiotemporal distribution of ion transporter proteins in the EO is essential for understanding electrocyte differentiation.

## Results

### Morphological observations of electrocytes

In *Electrophoru*, the tail region contains the mEO, which is the largest of the three organs responsible for electric discharge. In this study, especially in the histological analysis, the specimens were regarded as too young to observe well-developed Hunter’s and Sach’s organs; thus, they were excluded from the analysis. The mEO is located ventral to the skeletal muscle and consists of multinucleated syncytia called electrocytes (Figure 1A, 1B). The electrocytes were layered along the dorsal–ventral and anterior–posterior axes (Figure 1B). The series of cells along the anterior–posterior axis increases the membrane potential and generates a high-voltage discharge. Cell morphology along the dorsal–ventral axis is critical for understanding how electrocytes develop in the body; thus, we examined the histological morphology of electrocytes using Masson’s trichrome (MT) staining (Figure 1C, Figure S1). In the dorsal region of the mEO, we observed the typical structure of mature multinucleated electrocytes placed in a non-cellular compartment with an extracellular space around the cell body, which is important for generating a membrane potential for discharge. In the ventral region, such distinct extracellular spaces were not observed, and square cell bodies without protrusions filled the compartment (Figure 1D, Figure S1, S2). This morphological difference was also confirmed by the observation of toluidine blue-stained ultrathin sections and electron microscopy. Electron microscopy indicated that ventral electrocyte nuclei exhibited various shapes compared with those in the dorsal region. The presence and morphology of the mitochondria and caveolae showed no distinct differences between cells in the dorsal and ventral regions (Figure 1E). Through these experiments, we observed regional differences in electrocyte morphologies along the dorso-ventral axis.

**Figure 1.**
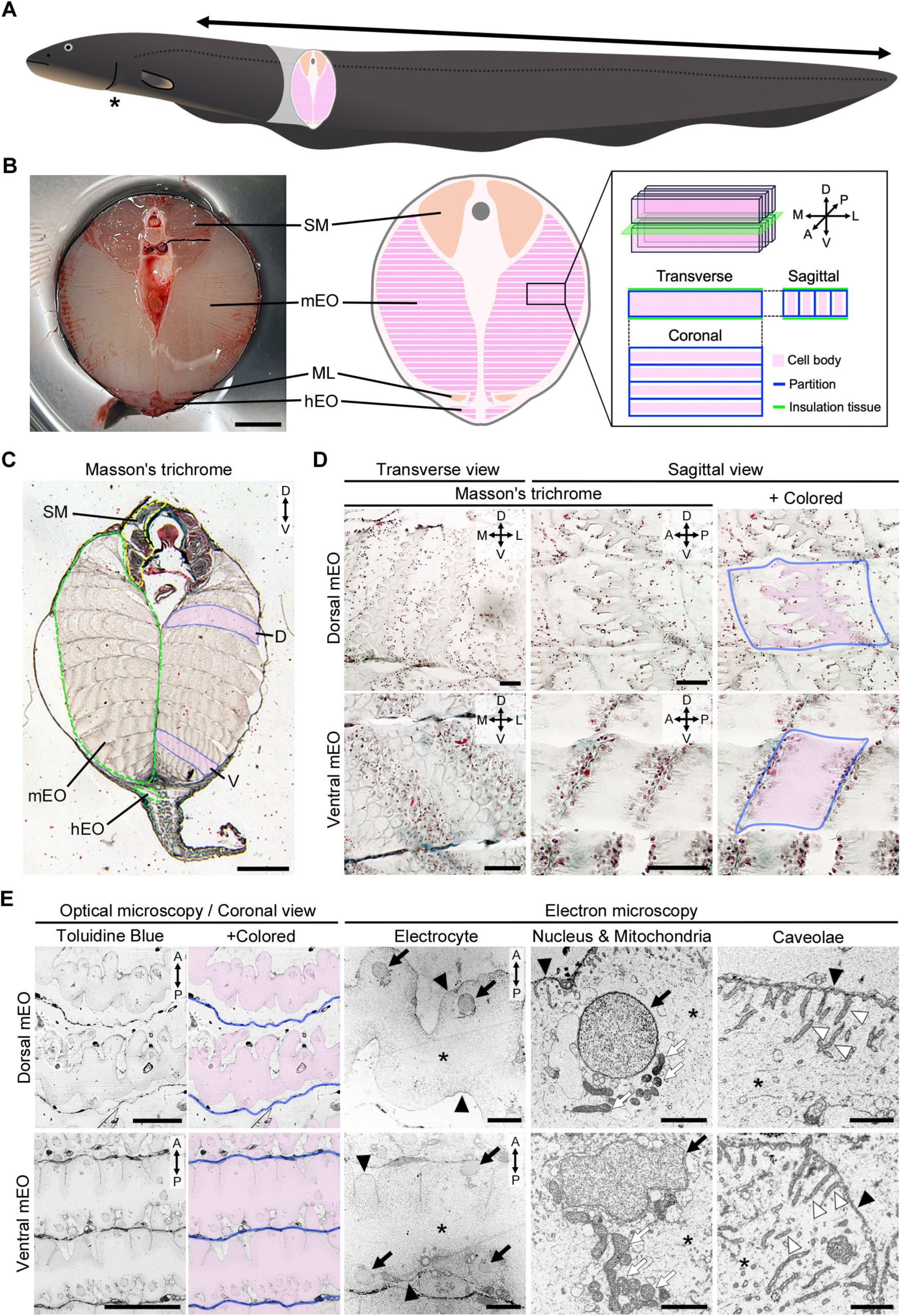
Morphological differences of multinucleated electrocytes. **A**. Illustration of the whole body of *Electrophorus*. The double-headed arrow corresponds to the tail region of the body. The asterisk indicates the anus. **B**. A photograph and illustration of the transverse section of the tail region of *Electrophorus*. The medial–lateral lines in the mEO indicate the outer edge of the multinucleated electrocytes. The plate-shaped electrocytes are stacked along the anterior–posterior and dorsal–ventral axes. SM, skeletal muscle. mEO, main electric organ. Scale bar, 1 cm. **C**. A whole transverse section of the tail region, including the electric organ. Scale bar, 1 mm. **D**. Cell morphologies of the electrocytes in the dorsal and ventral regions of the mEO. The blue lines indicate the outer edges of the electric unit. The cell bodies of electrocytes (colored magenta) are within the unit. Scale bars, 50 µm. **E**. Ultrathin sections (100 nm) of the electrocytes observed by optical or electron microscopy. The blue lines indicate the outer edges of the electric unit. The cell bodies of electrocytes (colored magenta) are within the unit. Black arrows, nucleus. Black arrowheads, plasma membrane. White arrows, mitochondria. White arrowheads, caveolae. Asterisks, cytoplasm. Scale bars, 50 µm (toluidine blue), 10 µm (electrocyte), 2 µm (nucleus & mitochondria), 1 µm (caveolae). A, anterior; D, dorsal; M, medial; L, lateral; P, posterior; V, ventral; hEO, Hunter’s electric organ; mEO, main electric organ; SM, skeletal muscle.

### Ventral terminal cell cluster in EO

The cell morphology of the mEO differed between the dorsal and ventral regions. Thus, a detailed examination of the cell morphology from the ventral to the dorsal terminus of the mEO was performed using HE and MT staining (Figure 2A, Figure S3, S4). A characteristic cell cluster consisting of high-density nuclei was identified at the ventral terminus of the mEO. This cluster was separated from the adjacent muscle layer by collagen fibers. In contrast, a structural border was not observed between the cluster and the dorsal region of the mEO (Figure 2B). The anti-myosin signal, which labels myogenic cells, was not detected in ventral cluster cells (Figure 2C, Figure S5). This indicates that the cluster belongs to the mEO, but does not possess the muscle lineage character. We then categorized the mEO into four populations based on cell and tissue morphology along the dorsoventral axis (Figure 2D, Figure S6). The terminal cluster consisted of high-density mononuclear cells located in the ventral terminus of the mEO, and the subterminal cell clusters were located in the dorsal position of the terminal cluster. The thin layers consisting of high-density nuclei appeared near the dorsal region. The thickness of the layers increased in the dorsal region, forming huge syncytial electrocytes (Figure 2E, Figure S7). Thus, the morphology of the cells in the mEO changed drastically along the dorsal–ventral axis. Fluorescent phalloidin labeling indicated that actin filaments were distributed in the ventral clusters and layers of the mEO (Figure 2F). Mono- or binuclear cells were observed at the edge of the layers in the ventral region. Immunohistochemistry for PCNA indicated that some of these cells, including binuclear cells, were undergoing the proliferation phase (Figure 2G).

**Figure 2.**
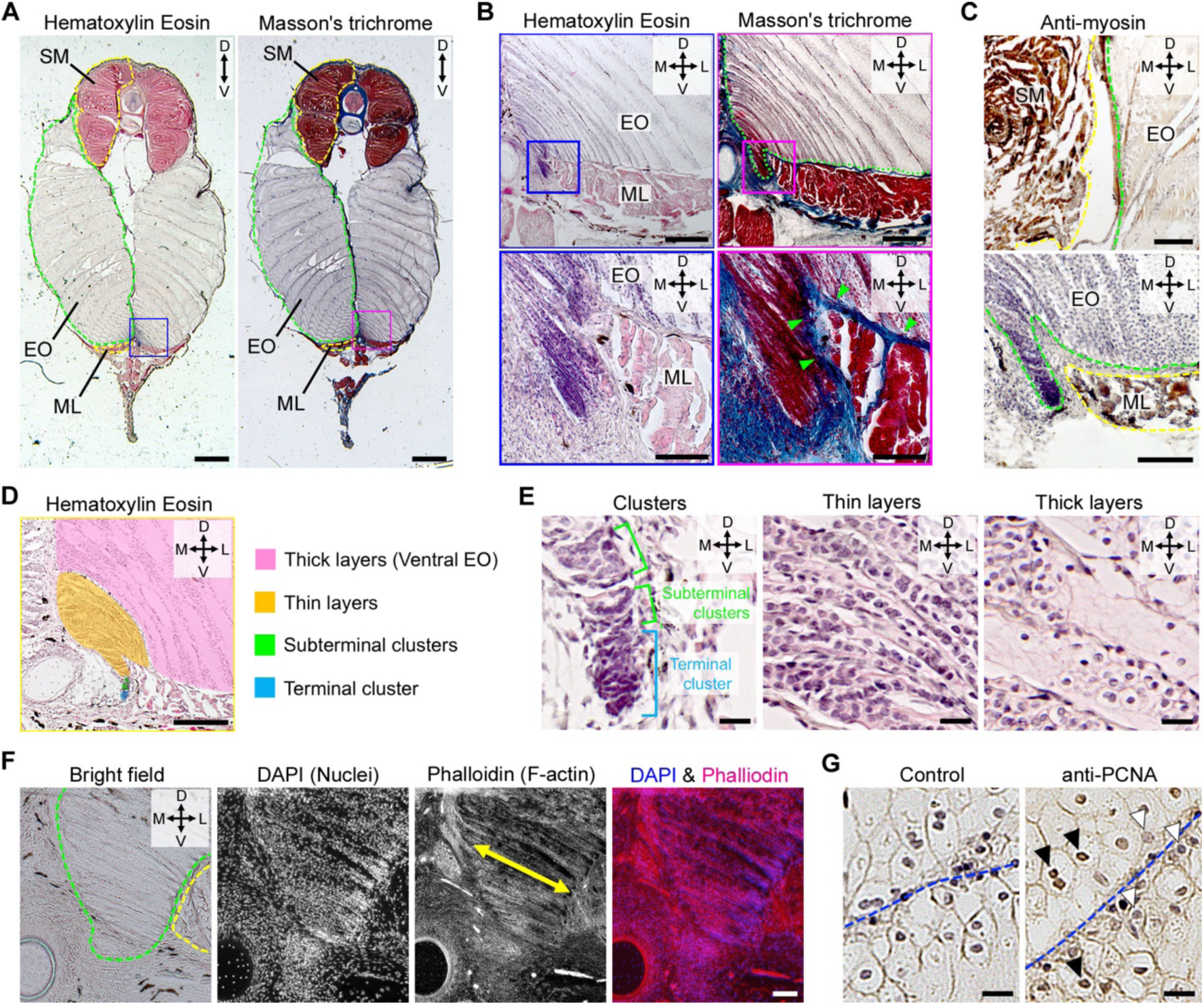
Morphological differences of cells along the dorsal–ventral axis. **A**. A whole transverse section of the tail region, including the electric organ. The green dotted line indicates the outer edge of the mEO. The yellow dotted lines indicate the outer edge of the muscular tissues. Scale bar, 1 mm. **B**. The ventral region of the mEO and the enlarged images of the ventral terminal cluster. The green dotted line indicates the outer edge of the mEO. The green arrowheads indicate the collagen fibers stained by Aniline Blue. Scale bars, 200 µm (low magnification), 50 µm (enlarged). **C**. Immunohistochemistry against myosin heavy chain in the dorsal and ventral positions. The muscular tissues show positive signals, but not in the mEO tissues. Scale bars, 100 µm (dorsal), 50 µm (ventral). **D**. A ventral region of mEO colored roughly by tissue structures. Scale bars, 100 µm. **E**. Enlarged images of each tissue structure in the mEO. The blue bracket indicates the terminal ventral cluster. The green brackets indicate the subterminal clusters. Scale bar, 10 µm. **F**. Fluorescent images of a ventral region of mEO with DAPI and phalloidin. The green dotted line indicates the ventral edge of the mEO. The yellow dotted line indicates the ventral muscle layer. The yellow double-headed arrow indicates the lateral width of the layer of the electrocytes. Scale bar, 50 µm. **G**. Immunohistochemistry against PCNA at the edge of the electrocyte layers. The blue dotted line indicates the boundary of the layer. The black arrowheads indicate the PCNA-positive nuclei. The white arrowheads indicate PCNA-negative nuclei. Scale bar, 10 µm. D, dorsal; M, medial; L, lateral; V, ventral; mEO, main electric organ; ML, muscle layer; SM, skeletal muscle.

Electron microscopy revealed the fine structures of the ventral region of the mEO (Figure 3A). The terminal cluster consisted of high-density mononuclear cells with scant cytoplasm (Figure 3B). Syncytial multinuclear cells with abundant cytoplasm were observed in the subterminal clusters (Figure 3A). Monocellular cells, similar to those in the terminal cluster, were also observed around the syncytium (Figure 3C). Intracellular components such as multiple nuclei, mitochondria, and actin filaments were observed in the syncytium (Figure 3D). Actin filaments were also visualized by phalloidin labeling (Figure 2F). The EO components were separated from adjacent muscles by collagen fibers (Figure 3E), and the presence of collagen fibers was also suggested by MT staining (Figure 2B).

**Figure 3.**
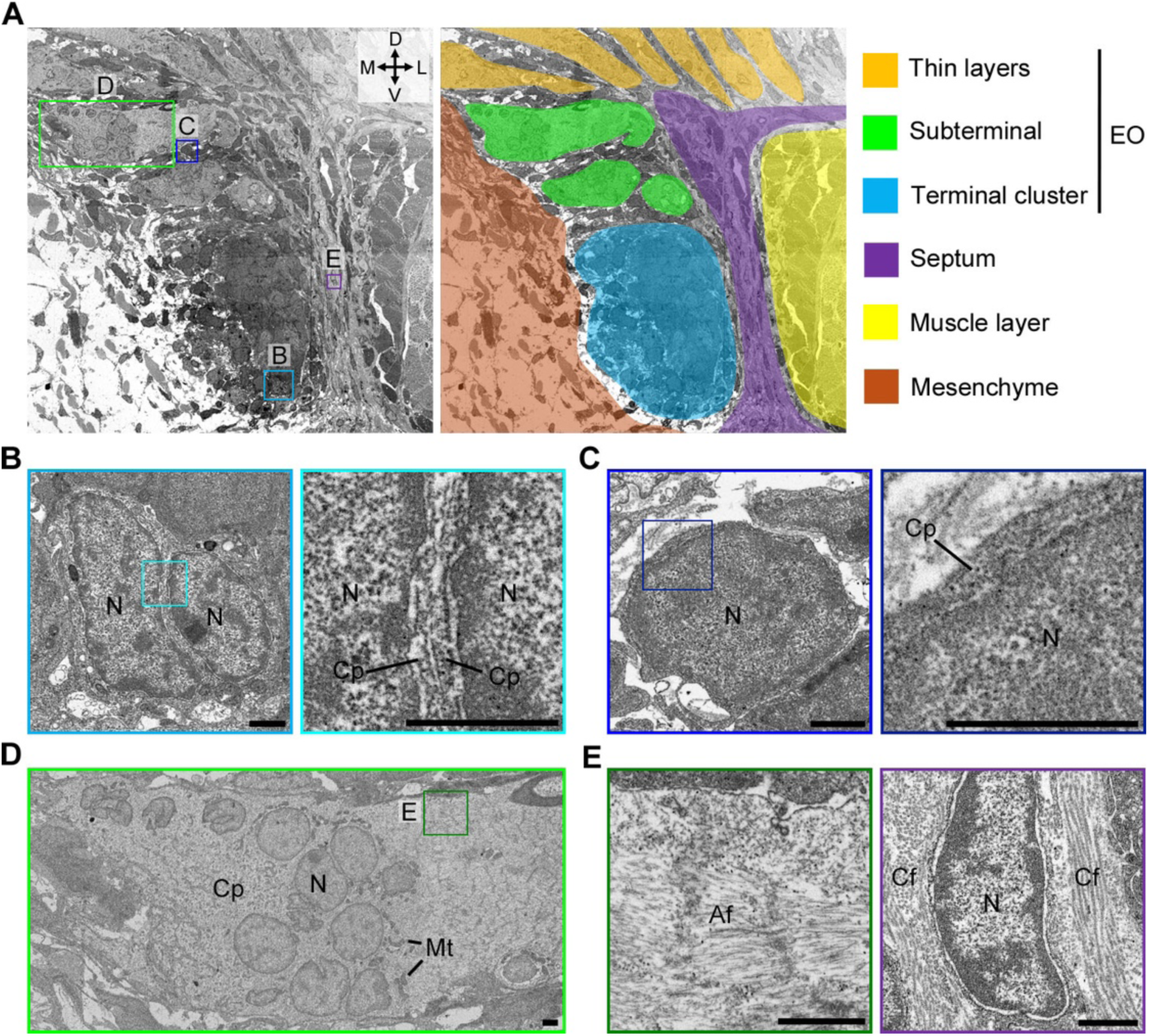
Fine structure of the ventral terminal region of the electric organ. **A**. Electron microscopy image of the ventral terminal cluster. The squares indicate regions of interest shown in Figures 3B–3E. The right panel presents the same region with rough coloring to highlight tissue structures. **B**. Two mononuclear cells with scant cytoplasm in the terminal cluster. The right panel outlines the cell borders. Scale bars, 1 µm. **C**. A mononuclear cell with scant cytoplasm adjacent to the syncytium in the subterminal region. The right panel highlights the cell’s edge. Scale bars, 1 µm. **D**. Intracellular structure of the multinuclear syncytium in the subterminal region. Scale bars, 1 µm. **E**. Fibrous structures in the syncytium (left) and the septum between the mEO and the muscle layer (right). Based on the histological labeling in Figure 2, the left structure corresponds to actin fibers, while the right corresponds to collagen fibers. Scale bars, 1 µm. D, dorsal; M, medial; L, lateral; V, ventral; Af, actin fiber; Cp, cytoplasm; Cf, collagen fiber; Mt, mitochondria; N, nucleus.

### Gene expression in EO

Tissue fragments from the mEO and skeletal muscles were excised via macroscopic surgery, and total RNA was extracted from each tissue. The RNA samples were subjected to next-generation sequencing (NGS), and the resulting sequence reads were processed to determine gene expression values and visualize differences in gene expression between tissues (Figure 4A and B, Table S1, S2). Our analysis revealed higher expression of genes in the mEO, including trophoblast glycoprotein like (*tpbgl*), N-myc downstream-regulated gene-4 (*ndrg4*), and others (Figure 4C and Table S3). From this dataset, specific protein groups with presumable functions—such as membrane transporters, scaffold proteins, and transcription factors—were selected, and their transcripts per million (TPM) values were compared between muscle and mEO (Figure 4D). This study focused on membrane molecules responsible for generating membrane potential through ion transport, such as acetylcholine receptors and sodium/potassium transporters, which exhibit higher expression levels in the mEO than in skeletal muscle. Notably, genes encoding sodium potassium adenosine triphosphatase (Na^+^/K^+^-ATPase, NKA) subunits (*atp1a2a* and *atp1b2b*) were highly expressed in the mEO, based on RNA sequencing. Cation transporters are believed to be localized in the anterior part of electrocytes (Figure 4E). In the tail region of *Electrophorus*, the electrocytes aligned along the anterior−posterior axis of the trunk, which is responsible for generating a high-voltage current due to synchronous activation regulated by motor neurons (Figure 4F).

**Figure 4.**
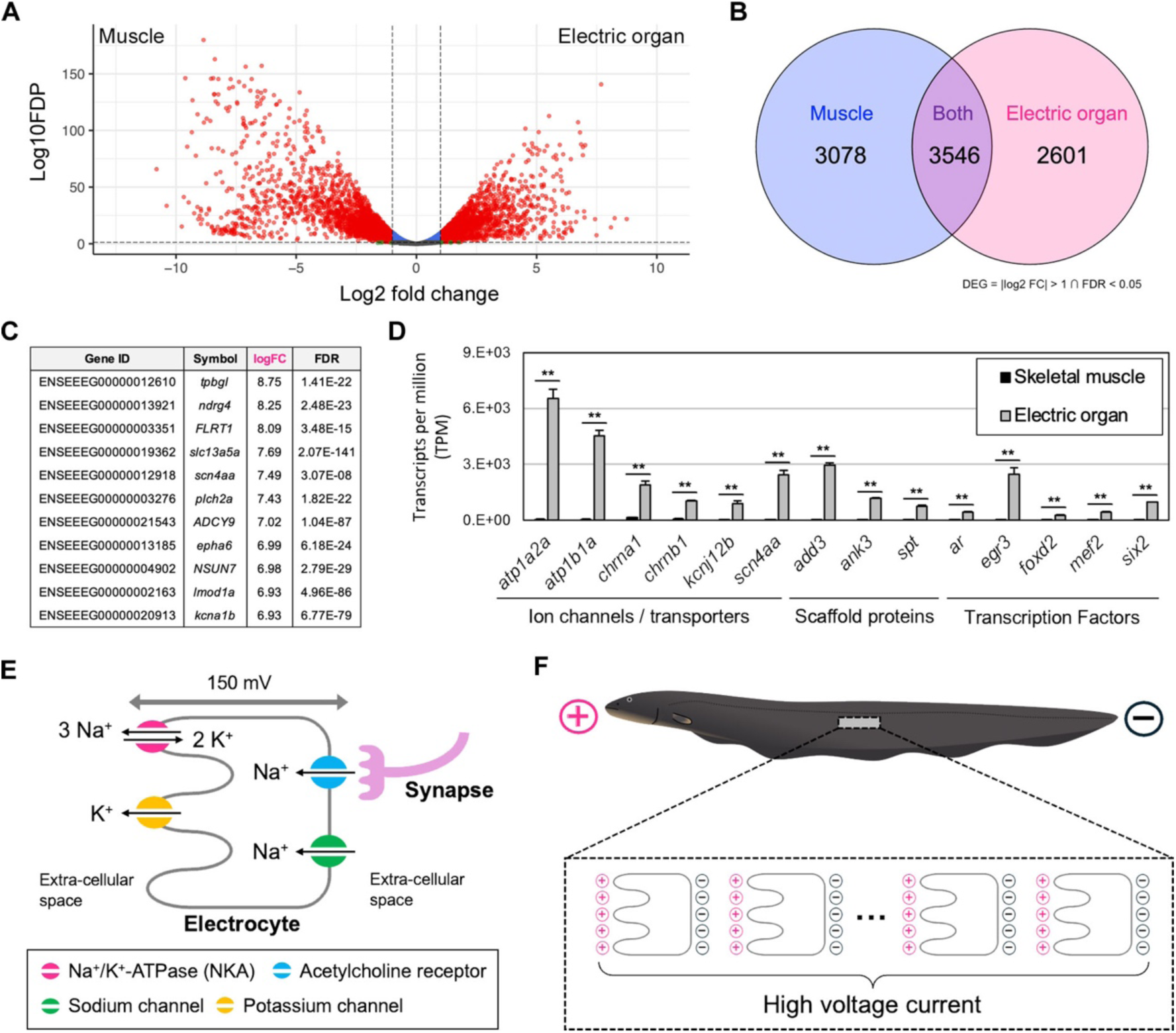
Gene expression and functions of ion transporters in electric organ. **A**. Volcano plot showing differentially expressed genes between skeletal muscle and mEO. Source data are deposited in the DDBJ (ID: PRJDB17973). **B**. Venn diagram illustrating differentially expressed genes in the mEO and skeletal muscle. **C**. List of differentially expressed genes sorted by logFC. **D**. Comparison of gene expression values (TPM) for selected genes differentially expressed in the mEO. **E**. Schematic model illustrating the generation of membrane potential in an electrocyte, with ion transporters localized anteriorly or posteriorly on the cell surface, generating approximately 150 mV current due to synaptic input. **F**. Electrocytes arranged along the anterior–posterior axis, enhancing membrane potential leading to high-voltage discharge.

### NKA distribution in EO

To evaluate electrocyte function, the presence and distribution of NKA protein in the mEO were detected using immunohistochemistry. The anti-NKA supernatant exhibited specific signals in the mEO and other organs, including the skeletal muscle and central nervous system, in *Electrophorus* (Figure 5A, Figure S8, S9). In the electrocytes, the plasma membrane of the mEO was labeled with the supernatant. The sagittal section indicated the accumulation of NKA protein in the anterior part of the plasma membrane (Figure 5B, Figure S8–S11). Additionally, the signals were lower in the ventral region of the mEO and absent in the ventral terminal cluster, even after longer chromogenic reactions (Figure 5C, Figure S12, S13). The NKA signals were not detected in the terminal cluster, even under conditions of prolonged incubation for DAB staining (Figure 5D). The border of NKA expression was observed in the thin layer, but the signals did not exhibit membrane localization, unlike in the thick layer, which includes the ventral EO (Figure 5E). Membrane localization of NKA signals was also observed in mono- or binuclear proliferating cells at the edge of the EO layers (Figure 5F). These data indicate that NKA is absent in the terminal cluster cells of the mEO but is upregulated and membrane-localized in the dorsal population cells, including mature electrocytes.

**Figure 5.**
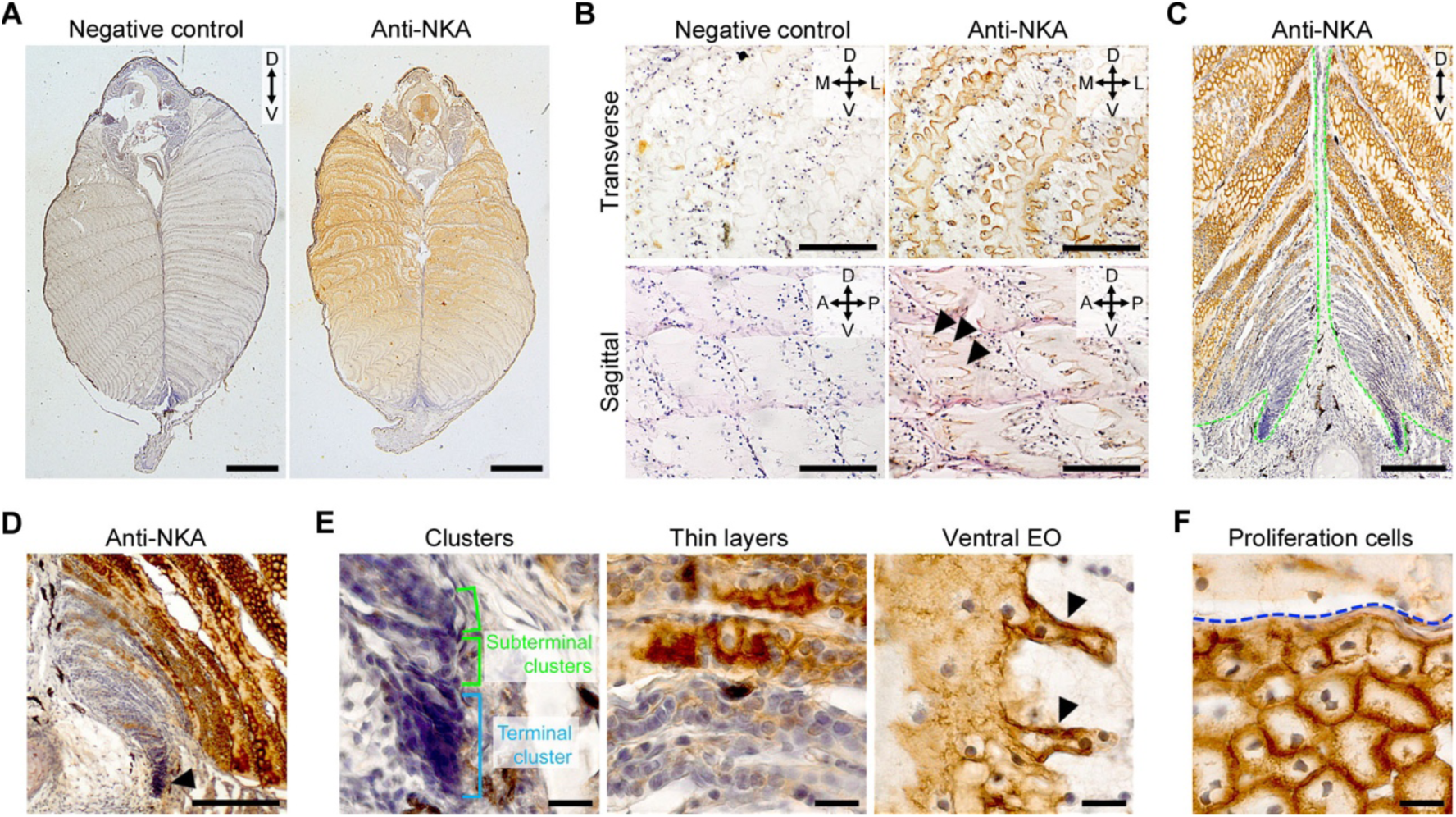
Presence and distribution of Na+/K+-ATPase in electric organ. **A**. Immunohistochemistry against NKA in transverse section. Scale bars, 1 mm. **B**. Enlarged images of anti-NKA immunohistochemistry in dorsal mEO in transverse and sagittal sections. Green arrowheads indicate anti-NKA signals on the anterior cell surface. Scale bars, 100 µm (transverse), 50 µm (sagittal). **C**. Ventral half of mEO in anti-NKA immunohistochemistry. Weaker anti-NKA signals observed in ventral terminal compared to dorsal regions. Scale bar, 100 µm. **D**. Immunohistochemistry against NKA in the ventral region of the mEO with prolonged incubation for DAB staining. DAB coloring was not detected in the terminal cluster (black arrowhead). Scale bar, 100 µm. **E**. Enlarged images of immunohistochemistry against NKA in different tissue structures of the mEO. The blue bracket indicates the terminal ventral cluster, while the green brackets indicate the subterminal clusters. Black arrowheads mark anti-NKA signals in the plasma membrane. Scale bar, 10 µm. **F**. Immunohistochemistry against NKA at the edge of the electrocyte layers. The blue dotted line indicates the boundary of the layer. Scale bar, 10 µm. A, anterior; D, dorsal; M, medial; L, lateral; P, posterior; V, ventral.

## Discussion

We analyzed the generation of electrocytes in *Electrophorus* using histology and immunohistochemistry. Our analysis suggested that ventral cluster cells are promising electrocyte progenitor candidates in the mEO at the juvenile stages. We propose that these progenitor cells differentiate into electrocytes and migrate from the ventral to the dorsal region. A previous study indicated that the first electrocyte at the embryonic stages is generated from the electromatrix located in the ventral region of the myotome (Schwassmann et al., 2014). However, nobody has demonstrated how the electric organ grows in the later stages; whether the electromatrix remains, or whether another source of electrocytes appears and functions. Our observation might provide an answer to these questions. In the juvenile stages, clusters of mononuclear cells were observed in the ventral terminal of the mEO, which is histologically similar to the electromatrix. To understand how electrocytes differentiate and how EOs grow, investigating the morphological and genetic characteristics of the ventral cluster cells is important.

Early studies alleged that EOs in electric fish were derived from muscle fibers or muscle lineage cells based on morphological observations. Histological observations of electric rays and larval mormyrid fish suggest that their electrocytes differentiate from mature muscle fibers (Fox and Richardson, 1978; Srivastava and Szabo, 1972). In *Electrophorus*, electrocytes were considered to differentiate from muscle based on light microscopy observations (Fritsch, 1883; Szabo, 1966). These studies concluded that the presumed electrocyte progenitors exhibited striated muscle-like structures. In our study, such muscular structures were not found in the ventral region of the mEO, and the ventral muscle layer was separated from the mEO by collagen fibers. Furthermore, the mEO lacked myosin-positive cells, which are characteristic of muscle tissue. Therefore, our results do not support the hypothesis that the electrocytes of *Electrophorus* are derived from muscle lineage cells. Other groups have claimed that electrocytes do not originate from differentiated muscle lineage cells but are directly developed from mesodermal cells before cell fate commitment (Keynes 1961). We do not have a definitive answer as to whether the terminal cluster cells are derived from the mesodermal lineage. Electron microscopy indicated that the terminal cluster is separated from the lateral muscle layer by collagen fibers but faces the medial mesenchymal tissue. To determine the origin of the electrocytes, further analyses investigating their cell lineage or gene expression profiles are required.

We identified a ventral terminal cell cluster in the mEO and observed continuous changes in cell morphology. Based on the clear histological images, we propose that the electrocytes originate as mononuclear progenitor cells in the ventral terminal cluster of the mEO and differentiate into multinucleated syncytia with dorsal migration. Our analysis indicated three findings: (1) morphological differences in electrocytes and their progenitors along the dorsoventral axis, (2) NKA expression and cellular distribution during the electrocyte development, and (3) timing of the syncytium formation (Figure 6). Although this study was limited by breeding methodology challenges in obtaining embryos under laboratory conditions and did not focus on embryonic stages, our findings on the source and differentiation of electrocytes during growth are crucial for understanding the molecular mechanisms underlying EO formation in *Electrophorus*.

**Figure 6.**
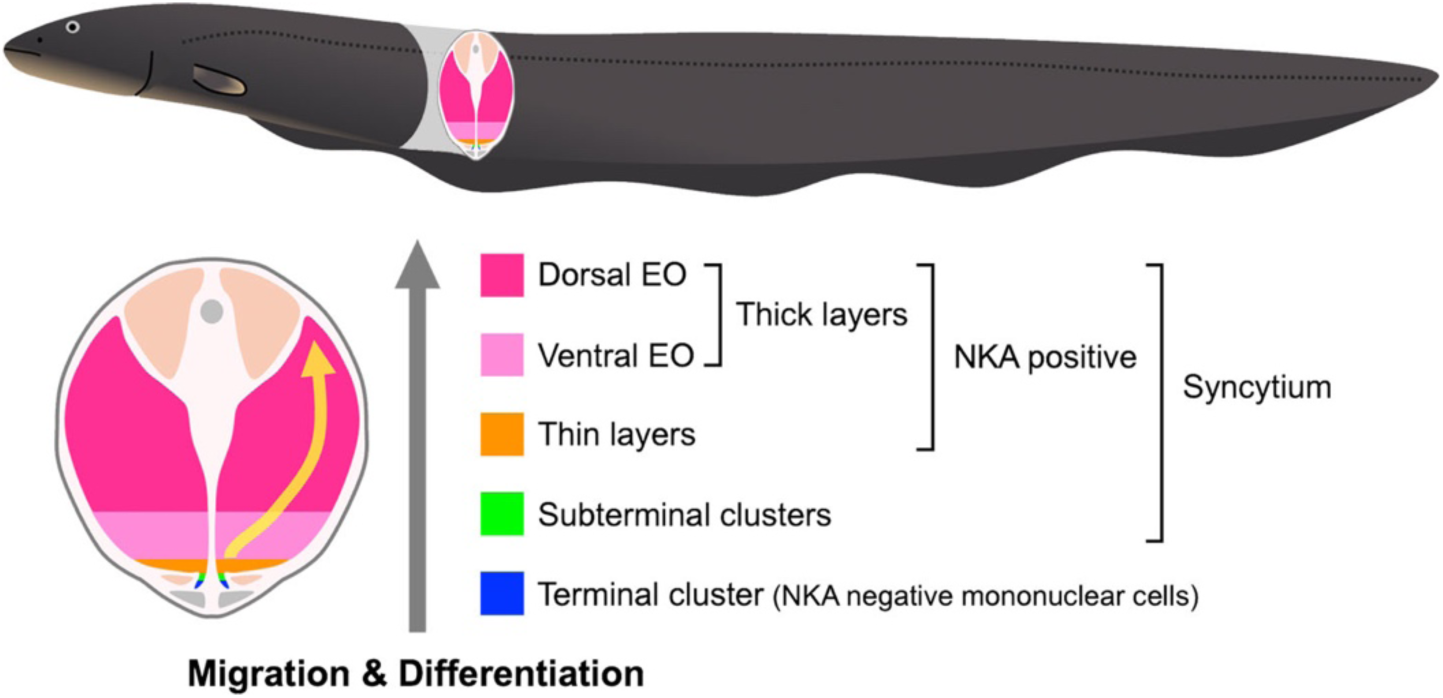
Model of main electric organ development. This illustration indicates the development of the main electric organ in the adult electric eel. Progenitor cells located in the ventral terminus of the main electric organ segregate and migrate dorsally, forming multinuclear syncytium and layer structure. The NKA ion pump begins to be expressed after the layer formation. The syncytia increase the dorsoventral thickness, differentiate into mature electrocytes, and contribute to the generation of high-voltage discharges via ion transporters, including NKA.

We focused on NKA as a functional molecule involved in generating membrane potential in electrocytes. We observed that the TPM values for the NKA subunits in the mEO were over 100-fold higher than those in the skeletal muscle. A previous study identified novel phosphorylation sites in the NKA protein of electric eels that were predicted to contribute to the high-voltage discharge of EOs (Traeger et al., 2017). NKA is ubiquitously distributed in animal tissues and plays an important role in generating membrane potential for cellular functions beyond those of electrocytes (Suhail, 2010). NKA is a popular sodium pump distributed not only in EOs but also in various types of cells; thus, anti-NKA signals were detected not only in the mEO but also in the skeletal muscle, kidney, brain, and other organs of *Electrophorus*. Notably, the ventral region of the mEO lacked anti-NKA signals, suggesting that ventral cluster cells are immature and unable to generate membrane potentials via the NKA ion pump. Cells in the ventral terminal cluster exhibit characteristic morphologies, such as small size, minimal cytoplasm, and round shape, which are typical characteristics of quiescent stem cells, including hematopoietic stem cells and muscle satellite cells (de Morree & Rando, 2023; Pinho & Frenette, 2019; Siegel et al., 2011). Therefore, we propose that the ventral terminal cluster includes stem cells that differentiate into electrocytes. Furthermore, single-cell RNA sequencing or cell characteristic evaluation using stem cell markers contributes to a deeper understanding of the maintenance of terminal cluster cells and their differentiation into electrocytes.

Methodological investigations using *Electrophorus* should be improved in future research. Our finding that multinucleated cells in the dorsal region of the mEO more closely resemble the typical morphology of functional electrocytes than those in the ventral region supports the conventional theory that the ventral region includes electrocyte progenitors (Gotter et al., 1998). We propose that the EO develops from progenitor cells in the ventral region and then progresses to maturity with dorsal migration. This ‘Ventral-to-Dorsal’ model does not conflict with previous reports (Szabo, 1966; Schwassmann et al., 2014). Our study enhances the understanding of the developmental process of EOs in *Electrophorus*. Further analyses using cutting-edge technologies, including NGS, are required to understand the molecular mechanisms by which EOs are generated in *Electrophorus*. A recent study proposed that changes in sodium channel expression are important for the evolutionary acquisition of EOs in weakly electric species (LaPotin et al., 2022). Darwin’s question arose from the independent acquisition of EOs in several fish lineages. The present study provides detailed information on the origin and developmental process of EOs in the electric eel *Electrophorus*. Future application of such molecular methods to other electric fish may ultimately lead to an answer to Darwin’s question.

## Methods

### Animal experiments

This study was approved by the Ethics Review Board for Animal Experiments at Nagoya University (approval number: A240105-002). All experiments involving live animals were conducted in strict accordance with institutional guidelines to ensure humane treatment. Electric eels used in this study were purchased from Meito Suien Co., Ltd. (Nagakute, Japan). The specimens used for histology were too young (TL: 10–15 cm) to determine their distinct species and sex; thus, we treated them as *Electrophorus sp*. (*Electrophorus*) in this study. The specimen used for sequencing analysis was a female with a developed ovary and was presumed to be *E. electricus* based on genome information.

### Histology and histochemistry

The electric eels were euthanized using a rapid cooling shock to ensure that they were unresponsive to gill movements and external stimuli (Wilson et al., 2009). Tail region cross-sections (5 mm thick) were dissected and fixed in 4% paraformaldehyde solution for 24 h at room temperature for histological observations. Fixed samples were then dehydrated in a graded ethanol series, cleared in xylene, embedded in paraplast, and sectioned at 4 μm thickness using a Leica microtome (Leica, Germany) (Kongthong et al., 2023; Presnell and Schreibman, 2013; Suvarna et al., 2013). Three consecutive sections were stained with Harris’ hematoxylin and eosin (HE) to observe body organization and tissue structure. MT staining was used to assess muscular structure and collagen fibers. Tissue samples were examined, photographed using an Olympus BX53 microscope, and imaged using a DP25 digital camera (Olympus, Shinjuku, Japan) or a BZ-X800 microscope (Keyence, Osaka, Japan).

### Electron microscopy

The tail region of *Electrophorus* was sliced into 5 mm transverse sections and fixed in 2.5% glutaraldehyde, followed by washing with PBS. The fixed specimens were treated with 1% osmium tetroxide in PBS for 60 min and dehydrated in a graded ethanol series (50%, 70%, 80%, 90%, 95%, 99.5%, and 100%). Samples were treated with propylene oxide (KISHIDA CHEMICAL, Osaka, Japan) and embedded in epoxy resin (Nissin EM Quetol-812). The blocks were polymerized at 60°C for 48 h. Sections of 1 μm thickness were stained with toluidine blue and observed using a Nikon ECLIPSE LV100ND with a DP74 digital camera (Olympus) to determine the region of interest. Sections of 100 nm thickness were cut using a Leica ultramicrotome and stained with UranyLess (Micro to Nano) and 3% lead citrate (Micro to Nano). Backscattered electron images were obtained using an SU9000 field-emission scanning electron microscope (Hitachi, Tokyo, Japan) at 1 kV.

### Phalloidin staining

Phalloidin staining was performed on frozen sections by using Alexa Fluor 546-conjugated phalloidin (Thermo Fisher Scientific, USA). Specifically, fixed samples were subjected to decalcification treatment in 10% EDTA (pH 7) for three consecutive nights. Following decalcification, the samples were cryoprotected by immersing them in 30% sucrose solution and subsequently embedded in the Optimal Cutting Temperature (OCT) compound (Sakura Finetek Japan, Japan). Frozen sections were cut into 10 μm-thick slices using a cryostat. The sections were stained with a 40-fold diluted Alexa Fluor 546-conjugated phalloidin solution at room temperature for 60 min. After staining, the sections were thoroughly washed with PBS and mounted using VECTASHIELD Antifade Mounting Medium with DAPI (Vector Laboratories, USA), which simultaneously enabled nuclear staining. Microscopic observations were performed under an Axioscope 5 microscope (Zeiss, Germany), and images were captured with an Axiocam 503 color camera (Zeiss).

### RNA sequencing

Total RNA was extracted from the mEO and skeletal muscle of an adult female electric eel using an RNeasy Plus Mini kit (QIAGEN). NGS was performed by Macrogen Japan Corp. (Tokyo, Japan) using a NovaSeq6000 platform (Illumina, Inc., San Diego, CA, USA). Approximately 40 million 100 bp paired-end reads were obtained for each sample. NGS data were deposited in the DNA Data Bank of Japan (DDBJ; ID: PRJDB17973).

### Transcriptomic analysis

The calculation of differentially expressed genes was performed using the National Institute of Genetics’ supercomputer at the Research Organization of Information and Systems. The adaptor and noise sequences were removed using the fastp program version 0.23.2 (https://github.com/OpenGene/fastp) (Chen et al., 2018). The cleaned reads were mapped to the genome of *E. electricus* (GCA_003665695.2, https://asia.ensembl.org/Electrophorus_electricus/Info/Index/) using the HISAT2 program version 2.2.1 (https://daehwankimlab.github.io/hisat2/) (Kim et al., 2019). The number of mapped reads was calculated using the featureCounts program in the subread package version 2.0.6 (https://subread.sourceforge.net/) (Liao et al., 2013, 2014). Further calculations were performed using the R software on a personal computer. Differentially expressed genes were identified using the edgeR package version 3.42.4 (https://bioconductor.org/packages/release/bioc/html/edgeR.html) with the following thresholds: false discovery rate < 0.05, and fold change > 2.00 (Robinson et al., 2010). A volcano plot was generated using the enhanced volcano package version 1.20.0 (https://github.com/kevinblighe/EnhancedVolcano).

### Immunohistochemistry

Consecutive sections from the histological protocol were prepared on slides coated with glycerol-albumin (Mayer’s fixative). The slides were deparaffinized in xylene, hydrated in a graded ethanol series, washed in tap water, and treated with 0.5% Triton X-100/PBS for 30 min and 0.3% H_2_O_2_ for 10 min to inactivate endogenous peroxidase activity. The sections were blocked with Blocking One solution for 2 h at room temperature (03953-95, Nacalai Tesque, Kyoto, Japan). The sections were incubated overnight at 4°C with either anti-myosin heavy chain (F59-s, DSHB, IA, USA, 1:5 in Blocking-One solution), anti-NKA α1 (a5-s, DSHB, 1:5 in Blocking-One solution) hybridoma supernatants, or PCNA Polyclonal antibody (10205-2-AP, proteintech, IL, USA, 1:1000 in Blocking-One solution). This was followed by incubation with Horse Anti-Mouse IgG Antibody (H+L) Peroxidase (PI-2000, Vector Laboratories, Inc., Burlingame, CA, USA, 1:500 dilution in 0.1% Tween-20/PBS) or Goat Anti-Rabbit IgG Antibody (H+L), Peroxidase (PI-1000, Vector Laboratories, Inc., Burlingame, CA, USA, 1:500 dilution in 0.1% Tween-20/PBS) for 2 h at 4°C. Staining was performed by incubation with 3,3’-diaminobenzidine tetrahydrochloride (SK-4105, Vector Laboratories, Inc.) for 5 min at room temperature. Finally, the slides were dehydrated and mounted using Permount (SP15-100, Thermo Fisher Scientific). A control section without primary antibody was prepared to ensure the specificity of staining. NKA localization was examined and photographed using the same method used for histological observations.

## Acknowledgments

Computations were partially performed on the supercomputer of the National Institute of Genetics (NIG) at the Research Organization of Information and Systems (ROIS). Yutaka Hattori arranged complimentary rentals of the digital microscope. Akihiko Koga and Akira Kanamori provided us with helpful comments. We are grateful for the constructive discussions with members of Unique-Kai, an annual meeting for biologists investigating non-conventional experimental animals in Japan.

